# Microglia-secreted TNF-α affects differentiation efficiency and viability of pluripotent stem cell-derived human dopaminergic precursors

**DOI:** 10.1101/2022.01.11.475933

**Authors:** Shirley D. Wenker, Victoria Gradaschi, Carina Ferrari, Maria Isabel Farias, Corina Garcia, Juan Beauquis, Xianmin Zeng, Fernando J. Pitossi

## Abstract

Parkinson’s Disease is a neurodegenerative disorder characterized by the progressive loss of dopaminergic cells of the *substantia nigra pars compacta*. Even though successful transplantation of dopamine-producing cells into the striatum exhibits favourable effects in animal models and clinical trials; transplanted cell survival is low. Since every transplant elicits an inflammatory response which can affect cell survival and differentiation, we aimed to study *in vivo* and *in vitro* the impact of the pro-inflammatory environment on human dopaminergic precursors. We first observed that transplanted human dopaminergic precursors into the striatum of immunosuppressed rats elicited an early and sustained activation of astroglial and microglial cells after 15 days post-transplant. This long-lasting response was associated with Tumor necrosis factor alpha expression in microglial cells. *In vitro* conditioned media from activated BV2 microglial cells increased cell death, decreased Tyrosine hydroxylase -positive cells and induced morphological alterations on human neural stem cells-derived dopaminergic precursors at two differentiation stages: 19 days and 28 days. Those effects were ameliorated by inhibition of Tumor necrosis factor alpha, a cytokine which was previously detected *in vivo* and in conditioned media from activated BV-2 cells. Our results suggest that a pro-inflammatory environment is sustained after transplantation under immunosuppression, providing a window of opportunity to modify this response to increase transplant survival and differentiation. In addition, our data show that the microglia-derived pro-inflammatory microenvironment has a negative impact on survival and differentiation of dopaminergic precursors. Finally, Tumor necrosis factor alpha plays a key role in these effects, suggesting that this cytokine could be an interesting target to increase the efficacy of human dopaminergic precursors transplantation in Parkinson’s Disease.

## Introduction

Neurological disorders are one of the main causes of disability in the world. Among these disorders, Parkinson’s disease (PD), affects the second largest group of people and has the highest growth in incidence. From 1990 to 2015, the prevalence of PD and, therefore, disability and cause of death doubled, with a prevalence of 98 cases of PD per 100,000 individuals, representing a 15.7% increase (1).

PD is a neurodegenerative disorder, whose cardinal pathology is the loss of dopaminergic (DA) neurons in the *substantia nigra pars compacta*. Current treatments for PD provide symptomatic relief but have side effects in the long term and do not halt disease progression or regenerate DA cell loss (2). Cell replacement therapy has been proposed as an alternative strategy due to the fact that motor symptoms of PD are caused by the degeneration of a specific cell type, DA neurons. Therefore, grafts of DA precursors (DAp) could have the potential to replace the function of DAn loss and thus reduce the associated motor symptoms. Pre-clinical and clinical trials have provided proof of concept that the transplantation of DA neuroblasts in the striatum can alleviate parkinsonian symptoms (3, 4). DA neuroblasts and not mature DA neurons are used since the transplanted cells need to engraft in the host parenchyma in order to survive. Although the results from clinical trials have demonstrated that this approach is safe, its efficacy was variable and several undesirable side effects such as graft-induced dyskinesia were reported. Thus, standardization of several factors is crucial to optimize the efficiency of the treatment and try to prevent unwanted effects (3, 5). One important factor for the development of an effective therapy of cell replacement is the host-primary response related to the graft. The microenvironment generated by the host could have dramatic effects on the survival, differentiation and proliferation of the transplanted cells. In this context, microglia activation affects dramatically the host environment after a mechanical intervention to the central nervous system (CNS). Cellular transplantation in the CNS not only includes such a mechanical injury but also provides a plethora of inflammatory and immune stimuli to the transplantation site (6). Adaptive immune responses are usually prevented by immunosuppression treatments. However, innate immune responses should remain active after transplantation but scarce information is available on its characteristics, duration and functional effects on the transplant. In particular, the possible effects of microglia, the main effectors of the innate immune response in the brain, on the fate of the transplanted cells are poorly described. Microglial activation produces several pro-inflammatory factors with neurotoxic effects (i.e., Tumor necrosis factor alpha (TNF-α), Interleukin (IL)-1, IFN-gamma, Nitric oxide, and reactive oxygen species) that could compromise the viability and/or differentiation of DA precursors (7). Because grafting to the CNS inevitably causes activation of host’s microglia, understanding its functional effects on the viability and differentiation of DA precursors is important in order to detect potential molecular targets to improve the efficacy of cell therapy.

In this work, we found that sustained and increased microglial and astroglial response were observed 15 days post-transplantation even in the presence of a constant immunosuppressive treatment. On the contrary, the transplanted cells elicited only a marginal and transient infiltration of neutrophils. Then, we observed that *in vitro*, activated microglial-derived conditioned media diminished human DAp survival, differentiation and affected cell morphology. These effects were blocked by inhibiting TNF-α in the culture.

## Materials and methods

### Reagents

All chemicals used were of analytical grade. D-MEM, αMEM; GMEM, Neurobasal, B27, Geltrex, Acutase, penicillin/streptomycin; NEAA; β-mercaptoethanol and KSR were obtained from Gibco. GDNF and BDNF were from Peprotech. L-glutamine; Na pyruvate; dimethylsulfoxide; Mitomicym C; lipopolysaccharide and p-formaldehyde (PFA) were obtained from Sigma-Aldrich. TNF-α -ELISA Kit was obtained from BD. Triton X-100 and Tween 20 were from Merck. Fetal bovine serum (FBS), and were obtained from Internegocios SA (Argentina). Cyclosporine was from Novartis.

### Cell cultures

#### PA6 cells

Mouse stromal cell line PA6 (RIKEN BRC) were maintained and propagated αMEM, supplemented with 10% FBS and antibiotics (100 U/ml penicillin, 100 μg/ml streptomycin).

### Human Neural Stem Cells (hNSCs)

hNSCs-H14, kindly gifted by Dr Xianmin Zeng were propagated using Neurobasal medium supplemented with B27, 2 mM NEAA, 20 ng/mL of bFGF and antibiotics (100 U/ml penicillin, 100 μg/ml streptomycin) on Geltrex-coated dishes. Quality control for hNSCs populations were analyzed by immunofluorescences for Nestin and SOX-1.

### Dopaminergic differentiation

Generation of PA6 conditioned media (PA6-CM): PA6 cells were cultured, grown to 80% confluence, and then treated with Mitomycin C (0.01mg/ml; 2 hs). After 5 washes with PBS, PA6 cells were incubated with fresh PA6 culture medium for 16 hs. Then, PA6 maintence culture medium was replaced with ESD medium (GMEM with 10% KSR, 1× NEAA, 1× Na pyruvate, and 1× β-mercaptoethanol). PA6-CM was collected every 24 h during 1 week (8). DA differentiation was initiated by culturing hNSCs with PA6-CM on culture dishes coated with poly-L-ornithine (20 µg/mL) and laminin (10 µg/mL). After 14 days of differentiation, DA precursors (DA14) (150000 live cells/cm2) were transferred to 24-well plates. Cells were cultured in PA6-CM with BDNF (20 ng/mL) and GDNF (20 ng/mL) until day 28 (DA28) (8).

The different states of maturation from hNSCs cultures to dopaminergic differentiation were analyzed macroscopically and by immunofluorescent staining of differentiation markers such as TH, Foxa-2, TUJ1, GFAP. The characterization was carried out at two stages of the differentiation protocol: day 14 (DA14) and day 28 (DA28) (9).

*In vitro* treatments were carried out after 24 hs of incubation of DA14 cultures.

### BV2 microglial cells

Mouse BV2 microglial cell line was provided by Dr. Guillermo Giambartolomei (Hospital de Clínicas, Buenos Aires, Argentina). Cell line was maintained in 100 mm plastic tissue-culture dishes (GBO) containing D-MEM supplemented with 2 mM L-glutamine, antibiotics (100 U/ml penicillin, 100 μg/ml streptomycin), 10% FBS (10).

All cultures were developed at 37°C with 5% CO_2_ and 100% humidity. Media were replaced every 2 days and cells were split before they reached confluence.

### Animals

Adult male Wistar rats (Jackson Laboratory, Bar Harbor, ME, USA), bred for several generations in the Leloir Institute Foundation facility, were used in all of the experiments. The animals were housed under controlled temperature conditions (22±2°C), with food and water provided ad libitum and a 12:12 dark:light cycle with lights on at 08.00 h. All experimental procedures involving animals and their care were conducted in full compliance with NIH and internal Institute Foundation Leloir guidelines and were approved by the Institutional Review Board “Cuidado y Uso de Animales de Laboratorio (CICUAL-FIL)”. All of the animal groups were periodically monitored, indicating that the welfare of the animals was consistent with the standards of the ethical guidelines for animals.

### Cell transplantation

For stereotaxic injections, the animals were anaesthetized with ketamine chlorhydrate (80mg/kg) and xylazine (8mg/kg). Intrastriatal stereotaxic transplantation was conducted on cyclosporine A-immunosuppressed male rats (age: 8 weeks). The stereotaxic coordinates were: bregma +1.0 mm; lateral +3.0 mm; ventral −5 and –4.5 mm (11). After confirmation of the viability of the hDAp with Trypan blue vital stain, the concentration of the suspension was adjusted to 125000 cell/µL. About 250000 human DAp (viability: 81%) derived from hNSCs-H14 were transplanted into the left striatum by using a Hamilton syringe (12). 2 μl of the cell suspension was inoculated. The injection flow was 0.5μl/min. After injection, the cannula was held in place for 5 min, before being slowly retracted. The experimental animals did not show signs of ongoing disease. They presented with normal fur, activity, movement, and food consumption.

Cyclosporine A (Novartis, 15 mg/kg) was administrated daily (Intraperitoneal injection) until the end of the experiment, starting 2 days before cell transplantation (13).

Host primary response related to the graft were analysed by histological and immunofluorescences techniques at different time points (1, 7, 15 and 28 days. *n*=5 rats per group). In this case, the male rat was the experimental unit.

An aliquot of hDAp obtained from transplantation were cultured for terminal differentiation *in vitro.* At DA28, 17±1% of Tyrosine hydroxylase (TH)-positive cells were detected by immunofluorescence.

### *In vitro* experimental treatments

A) Microglia activation: BV2 cells were seeded in 6-wells plate (75000 cells/cm2). After 24 hs, cell cultures were treated with lipopolysaccharide (LPS) for 24 h according to Dai and collegues (10).

B) DA precursors acute exposure to conditioned media from microglial cells: DA precursors had been previously plated for 24 h, as DA14, in 0.5 ml of PA6 medium containing differentiation factors. Activation of BV2 microglial cells (24 h) were carried out as explained before. After centrifugation (2000 rpm, 10 minutes) 0.6 ml cell-free supernatants (CM, conditioned media) from basal and activated microglial cultures were transferred to wells containing DA precursors (DA15). After 4 days of DA precursor incubation with conditioned medium from BV2 cells (CM-BV2), evaluation of cell survival and differentiation was performed.

In TNF-α neutralization experiments etanercept (Enbrel, Pfizer; 100 ng/mL) (14), an inhibitor of both soluble and transmembrane forms of TNF, was added to CM-BV2 as co-treatment.

### Measurement of nitric oxide

Nitric oxide (NO) production was determined by measuring the accumulation of nitrite, the stable metabolite of NO, in culture medium. Isolated supernatants collected from microglial cell cultures exposed to LPS for the indicated period were mixed with equal volumes of the Griess reagent (1% sulfanilamide, 0,1% naphthylethylenediamine-dihydrochloride, and 2% phosphoric acid) and incubated at 25°C for 10min. Absorbance at 540 nm was measured in a microplate reader (15).

### Cytokine quantification

TNF-α was measured by enzyme-linked immunosorbent assay (ELISA) in supernatants obtained as described above for NO measurement. ELISA test was performed according to the manufacturer’s instructions of the kit. In each trial, samples were analyzed in duplicate against standards of known concentration.

### Histology

The animals were deeply anaesthetized as previously described (16) and were transcardially perfused with heparinized saline, followed by ice-cold 4% PFA in phosphate buffer (PB) (0,1M; pH 7,2). Brains were dissected and placed in the same fixative overnight at 4°C and cryoprotected in 30% sucrose 0.1M PB solution. Then, the brains were frozen in isopentane and cut using a cryostat into 40-μm serial coronal sections through the left prefrontal cortex. Sections were mounted on gelatine coated slides and stained with Cresyl Violet to assess the general nervous tissue integrity, neutrophils cell counts and inflammation. For immunohistochemistry, sections were stored in cryoprotective solution at −20°C until needed.

### Immunofluorescence

Free-floating sections were rinsed in 0.1% Triton in 0,1M PB. After washes with PB-T, samples were blocked in 1% donkey serum for 45 min, and then incubated overnight at 4°C with primary antibodies diluted in blocking solution. The list of antibodies is provided in Table 1. After three 10-minute washes with 0.1 mol/L PB, the sections were incubated with indocarbocyanine (Cy3) or cyanine Cy2 (Cy2)-conjugated donkey anti-rabbit or anti-mouse antibody, respectively (1:500; Jackson ImmunoResearch) for 2 h at room temperature, rinsed in PB and mounted in Mowiol (Calbiochem). Digital images were obtained in a Zeiss LSM 510 laser scanning confocal microscope equipped with a krypton-argon laser.

**Table 1:** list of antibodies used

| Antibody | Trademark | Catalogue numbers |
| --- | --- | --- |
| Anti-GFAP | DAKO | Z0334 |
| Anti-MHCII | Serotec | MCA46G |
| Anti-ED1 | Serotec | MAC341R |
| Anti-SOX1 | R&D system | AF3369 |
| Anti-Nestin | Abcam | ab22035 |
| Anti-TUJ1 | Promega | G712A |
| Anti-MAP-2 | Cell Signaling | 9043S |
| Anti-Foxa-2 | Abcam | ab117542 |
| Anti- Tyrosine hydroxylase (TH) | Pell-Frezze | P40101 |
| Anti-Caspase 3 active (CA3) | Neuromics | RA15046 |
| Anti-NfκB | SCBT | sc-372 |
| Anti-hNCAM | SCBT | sc-33686 |
| Anti-TNF-α | SCBT | sc-1351 |

DA cell cultures: samples were fixed with PFA in 4% w/v in PBS, with incubation at room temperature (20 min). After three washes with PBS-T, samples were blocked with 1% donkey serum in PBS-T (1h at room temperature), and then incubated with primary antibody (Table 1), for 16hs at 4°C. After PBS-T washes, incubation with the secondary antibody (1:1000; Jackson ImmunoResearch) were performed proceed for 2 hrs at room temperature in the dark. At the end, excess antibody was removed with three new washes with PBS, nuclear staining will be performed with Hoechst 33258 dye (1:1000 dilution in PBS). After washing thrice with PBS, samples were mounted in Mowiol (Calbiochem).

BV-2 cells: After fixation with methanol, NFκB immunocytochemistry was performed on BV2 cells to determine nuclear translocation as a proxy for cell activation. Cells attached to coverslips were permeabilized with 0.5% triton X-100 in PBS and unspecific binding sites were blocked with 1% BSA in PBS. Cells were incubated with primary antibody against NFκB and then with Alexa Fluor (488/596; Thermo-Fisher) labeled secondary antibodies. Nuclei were visualized Hoechst 33258 dye (1:1000). After washing thrice with PBS, samples were mounted with Mowiol (Calbiochem).

### Fluorescence microscopy: Hoechst nuclear staining of apoptotic cells

DA cell cultures were developed on slide covers plated in in 24-wells plate. After treatments, cells were fixed with PFA 4% w/v in PBS for 20min at 4 °C, exposed to Hoechst 33258 dye in PBS for 30 min at room temperature, and washed thrice with PBS. Finally, samples were mounted and fluorescent nuclei with apoptotic characteristics were detected and analyzed by immunofluorescence microscopy. Apoptotic cells were identified by morphology and nuclear fluorescence intensity. The condensed chromatin within apoptotic cells stains particularly heavily showing blue fluorescense. In addition, little apoptotic bodies released from nuclei are also detected because of their brilliant blue color. Differential cell count was performed by evaluating at least 1000 cells (15).

Images were captured with Zeiss Axio Observer and Zeiss LSM 510 laser scanning confocal microscope equipped with a krypton-argon laser.

### Image analysis

Polymorphonuclear-neutrophil (PMN) cells were identified by their nuclear morphology appearance in 40-μm thick cresyl violet-stained sections. For MHC II-positive and GFAP-positive cell quantification, approximately 10 fields were quantified for each animal using the Zeiss Image J software. The total number of positive cells was normalized to the total area counted for each sample (16). Twenty images were obtained by random sample and the analysis was performed using the Image J software. Using the image overlay and cell count plugings, the total number of cells per field and the number of cells positive for the corresponding labeling were counted (example: TH, TUJ1, Foxa-2). In this way, the percentage of positive cells will be obtained in relation to the number of nuclei counted. At least 1000 total cells were counted.

### Differentiation analysis

DA precursors were seeded and after 24 h-incubation, DA15 were exposed to CM-BV2. Cell morphology was observed under phase-contrast in an inverted microscope at 200x magnification and photographed by using a Nikon DS-L3 digital camera. Cell differentiation was determined in independent experiments by counting differentiated cells based on morphological criteria (17) and expressed as percentage of total cells (at least 500 cells).

For neurite outgrowth analysis from TH+ cells derived from DA differentiation were performed at DA19 and DA28 stages. Multiple independent images were taken from immunofluorescence against TH at 400x magnification. At least 10 neurites per field were selected for neurite outgrowth measurement and means of neurite length were calculated for each assay. This neurite tracing technique was implemented in the form of a plugin (NeuronJ) for ImageJ, the computer-platform independent public domain image analysis program inspired by NIH-Image (18).

### Statistical analysis

Statistical analysis was performed using GraphPad Prism, version 6.00 for Windows (GraphPad Software, San Diego, CA, USA). Results were expressed as mean and standard error (SEM) of n independent trials (at least 3) indicated in each figure. Statistical significance was calculated using two-tailed Student’s t test or ANOVA followed by *post hoc* multiple comparison test as indicated. When corresponding statistical differences between groups were assessed by means of the Mann-Whitney test or the Kruskal-Wallis One Way Analysis of Variance. At least differences with P<0.05 were considered the criterion of statistical significance.

## Results

### Host response to hDAp transplantation

hNSCs expressed over 95% of Nestin and Sox-1, proving that they were *bona fide* hNSC (Fig. 1C). Human dopaminergic precursors (hDAp) were obtained by incubating hNSCs with PA6-conditioned medium (PA6-CM) (Fig.1A). We observed morphological changes such as outgrowth of elongated cells (Fig. 1B). After 14 days, DA precursors (DA14) expressed characteristic early DA markers such as Foxa-2 (72,6±4,7 %) and late markers such as Tyrosine hydroxylase (TH) (5,5±0,8 %). By 28 days, an increase in neuron-like cells was observed. As expected for such a protocol, at 28 days, cultures contained approximately 19,1±0,9% TH-positive cells; 70,5±7,3 % positive cells for the pan-neuronal marker, TUJ1; and less than 5 % of glial cells (3,4±0,5 % GFAP-positive cells) (Fig.1C).

**Fig 1.**
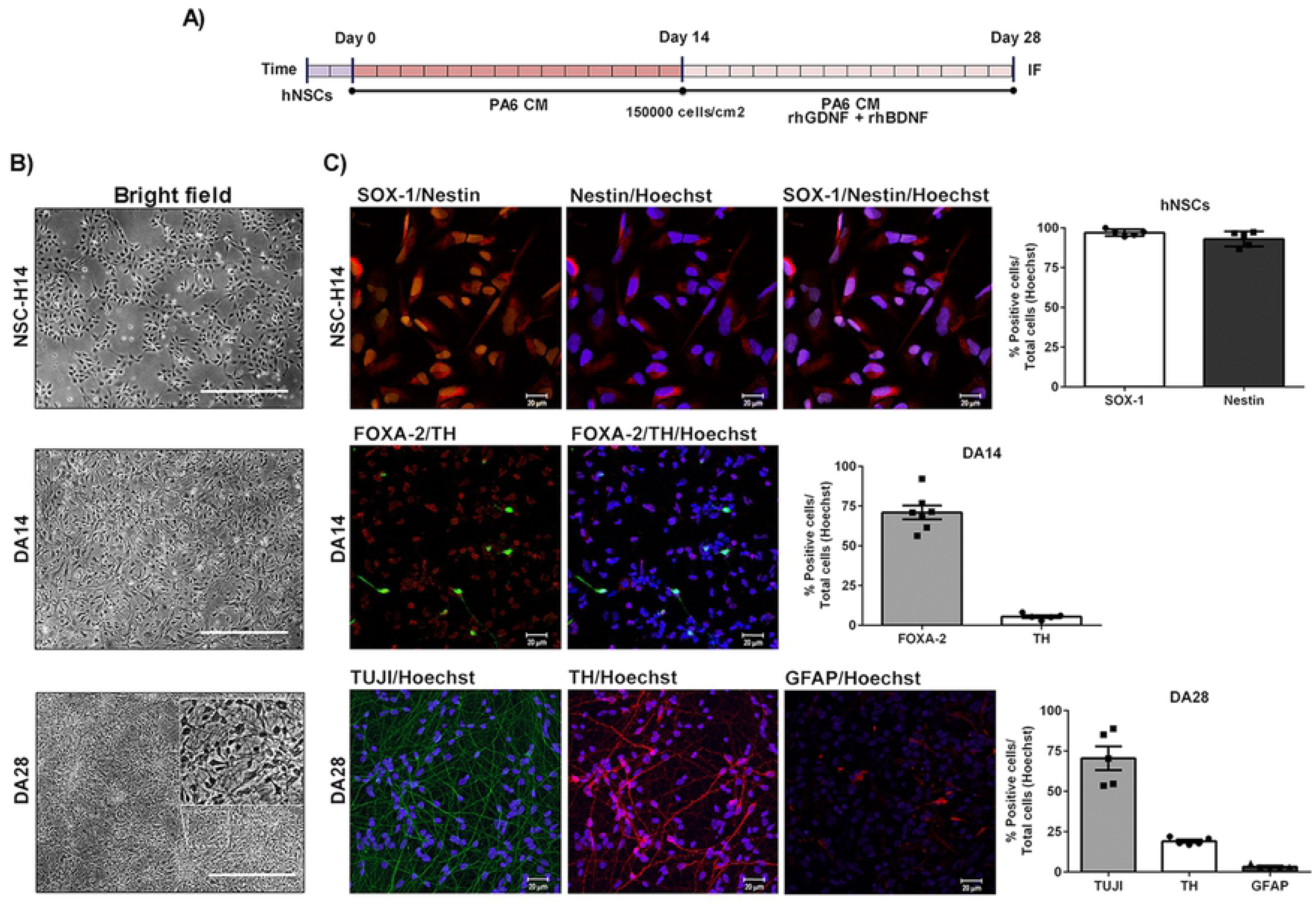
Dopaminergic differentiation from hNSCs. (A) Differentiation protocol was initiated by culturing hNSCs-H14, in PA6-CM for 14 days (DA14). After this period, PA6-CM was supplemented with rhBDNF and rhGDNF for a period of 14 days (DA28). (B). Photographs from each stage shown morphological changes. (C) Quality control for hNSCs populations were analyzed by immunofluorescence technique for Nestin and SOX-1. Different stages of maturation from cultures of hNSCs induced to dopaminergic differentiation were analyzed by immunofluorescence. The expression of the following markers were detected and quantified at two stages of the differentiation protocol: Foxa-2 and TH for DA14 cells (DA precursors) and TUJI; GFAP and TH for DA28. Cell nuclei were labelled by Hoechst staining. Each bar represents mean±SEM of independent assays. Magnification: 40X.

A preparation of 250.000 hDAp with at least 80% living cells was trasplanted into the striatum of cyclosporine A-daily immunosuppressed rats during 28 days (Fig. 2A) (12). Post-mortem evaluation revealed surviving grafts, visualised by the presence of human-specific NCAM staining (Fig.2B).

**Fig 2:**
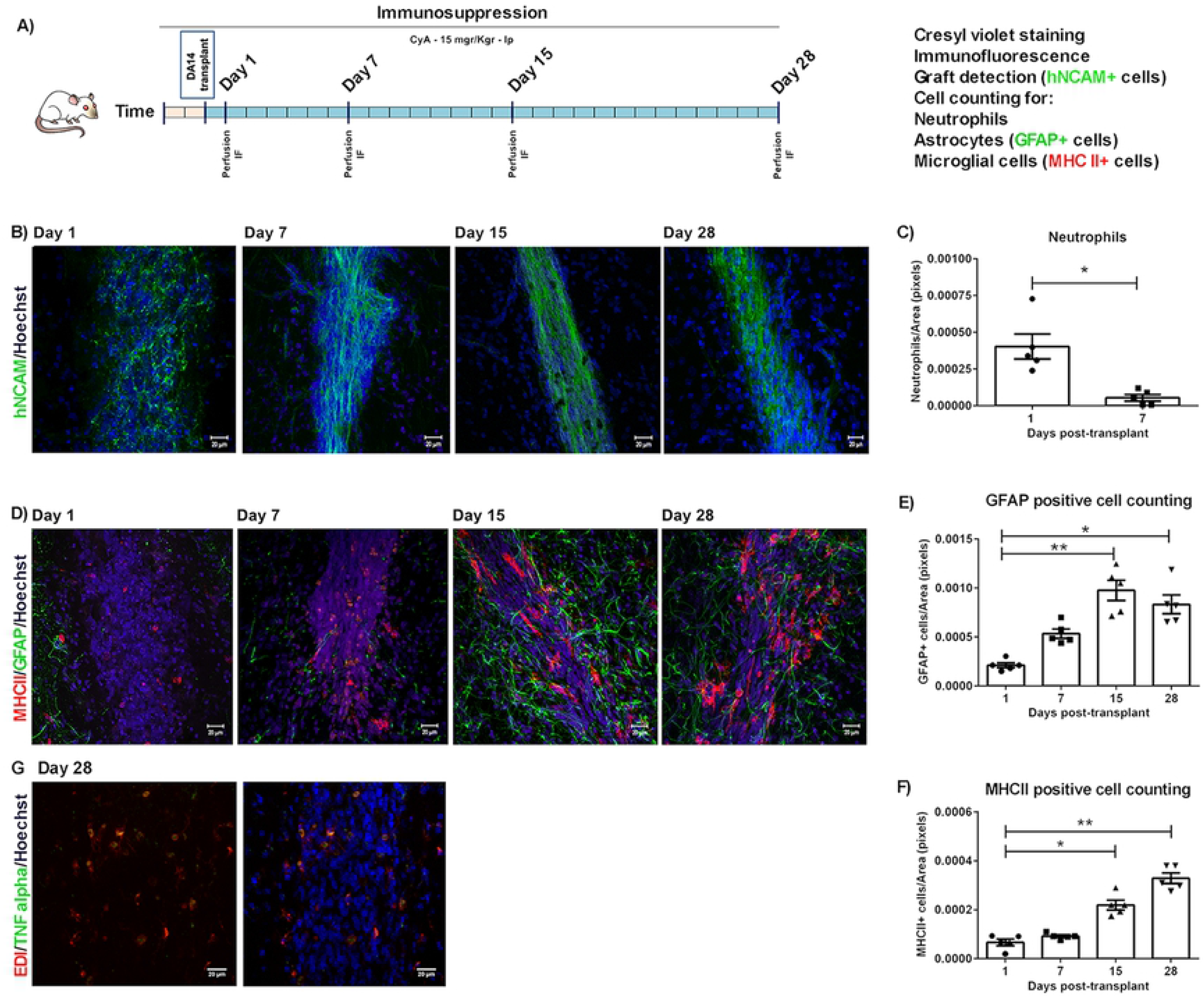
Primary host response related to the graft. (A) DA14 cells were obtained from hNSCs and 250000 cells (viability: 81%) were transplanted into the striatum of immunosuppressed Wistar male rats (age: 8 weeks). At different time points (1, 7, 15 and 28 days) host primary response related to the graft were analysed by immunofluorescences technique. (B) Detection of human-specific NCAM (hNCAM) allowed for identification of the graft. (D) Astrocytes (GFAP-positive cells) and activated microglia (MHCII-positive cells) related to the graft can be observed after 7 days of surgery. (E-F) There is a significant increase in MHCII+ and GFAP+ cells at 15 and 28 days post graftment of hDA14 (*p<0,05 and **P<0.01). Kruskal-Wallis ANOVA followed by Dunn’s post hoc test, n=5. (G) Finally expression of TNF-α were observed at Day 28 post surgery. Representative pictures of the grafts are shown (Magnification: 40X). Each bar represents mean±SEM of independent assays

Histological analysis revealed an early neutrophil infiltration (1day: 4,04e-4±8.50e-5 Neutrophils/area) from the periphery, which was resolved by 7 days (5,46e-5±2,25e-5 Neutrophils/area. *p<0,05 vs. 1 day) (Fig. 2C). Astroglial activation as seen by GFAP-staining, increased 7 days after transplantation, reaching a peak at 15 days (1 day: 2,13e-4±2,54e-5 vs. 15 days: 9,76e-4±1,03e-4 GFAP+cells/area. **p<0.01), beginning to decay at 28 days (28 days: 8,34e-4±9,58e-5 GFAP+cells/area. *p<0,05 vs. 1 day) (Fig. 2D, 2E). Transplantation of hDAp in cyclosporine A-daily immunosuppressed rats resulted in microglial activation as demonstrated by the presence of MHC-II+ cells, which specifically label activated microglia (16, 19). MHC-II stained all the activation stages but not resting microglia. According to our results, MHC-II+ cells presented the typical morphology of activated microglial cells: elongated-shaped cell body with long and thicker processes and/o round-shaped body with short, thick and stout processes (Fig. 2 D). Statistical analysis of microglial activation determined by MHCII-positive cell density showed a marked increase at 15 days (1 day: 6,67e-5±1,32e-5 vs. 15 days: 2,19e-4±1,96e-5 MHCII+cells/area. *p<0,05) which was sustained up to the last time point analysed 28 days: 3,29e-4±2,16e-5 MHCII+cells/area. **p<0.01 vs. 1 day) (Fig.2D, 2F).

Interestingly, tumor necrosis factor alpha (TNF-α) expression was detected 28 days post-surgery in ED1-positive microglial cells within the graft core and the periphery (Fig. 2G). These results show that there is a sustained host response to the transplant in immunosuppressed animals.

### Conditoned media from microglia have toxic effects on hDAp: short term effects

Previously, we observed a primary response related to the xenograft of hDAp, with a significantly increase of microglial activation. As the effects of the host microglial cells on the fate of the transplanted dopaminergic precursors are poorly described, we were interested in simulate the impact of rodent microglial activation on the differentiation and survival of hDAp derived from NSCs.

Microglial activation using bacterial lipopolysaccharide (LPS), was analysed by determining pro-inflammatory mediators such as nitrites (NO) and TNF-α in the cell culture supernatant. LPS led to a significant increase in nitrite production (Basal: 1,2±0,7 vs. Activated (LPS): 31,9±1,5 uM. **P<0.01) and TNF-α secretion (Basal: 22,3±14,3 vs. Activated (LPS): 739,7±76,1 pg/ml. *P<0.05) 24 h after cell activation (Fig. 3A, 3B). In addition, NFkB nuclear translocation was also observed in BV2 cell cultures after cell activation (Fig. 3C).

**Fig 3:**
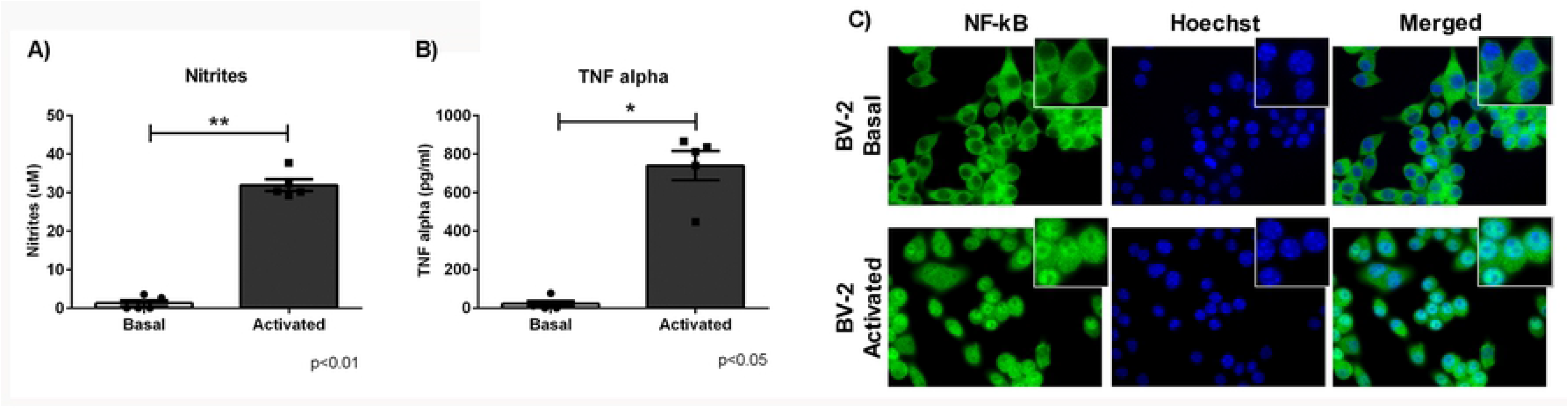
Characterization of microglial cell activation. BV2 cultures were exposed or not (Basal) to LPS (Activated), and parameters of microglial cell activation were determined. (A) Nitrite production (Griess method) and TNF-α secretion (ELISA test) (B) were measured after 24 h of cell activation. (C) NFkB nuclear translocation was analyses by immunofluorescences after short term exposure to LPS. Significant increases in nitrite and TNF-α were induced by LPS treatment. Asterisks indicate statistically significant differences: nitrite, **P<0.01 with respect to basal condition (n=5) and *P<0.05 TNF-α with respect to basal condition (n=5) (Mann-Whitney test). Microscopy images of NFκB-stained microglia under basal conditions or activated after LPS treatment (Magnification: 40X). Values are means±SEM of n independent trials.

In order to analyze the short-term effect of microglial activation on hDAp, we exposed DA15 cultures to conditioned medium from BV2 cells (basal –CM-BV2 Basal- or activated condition –CM-BV2 Activated-) until DA19 (Fig. 4A). Morphological analysis were performed. A decrease in neuron-like cell count (CM-BV2 Basal: 72,8±1,9 vs. CM-BV2 Activated: 57,9±2,4 %.***p<0,001. Figure 4B,4C) and neurite length of TH positive cells was detected in DA precursors cultures incubated with CM from activated microglia (CM-BV2 Basal: 339,0±31,2 vs. CM-BV2 Activated: 148,0±15,4 neurites lenght.***p<0,001) (Fig. 4B-E). A significant reduction of TH-immunoreactive cells was detected in DA cultures exposed to CM from activated BV2 compared to basal conditions (CM-BV2 Basal: 5,9±0,4 vs. CM-BV2 Activated: 2,5±0,4 %TH+cells. ***p<0,001. Fig. 4F, 4G). Concomitant with these results, we observed an increment in apoptotic nuclei (CM-BV2 Basal: 7,2±0,7 vs. CM-BV2 Activated: 21,4±2,5 % apoptotic cells. **p<0,01. Fig. 4I) and caspase activated 3 (CA3)-positive cells (Fig. 4H) in DA precursors after 4 days of incubation with CM from activated microglia.

**Fig 4.**
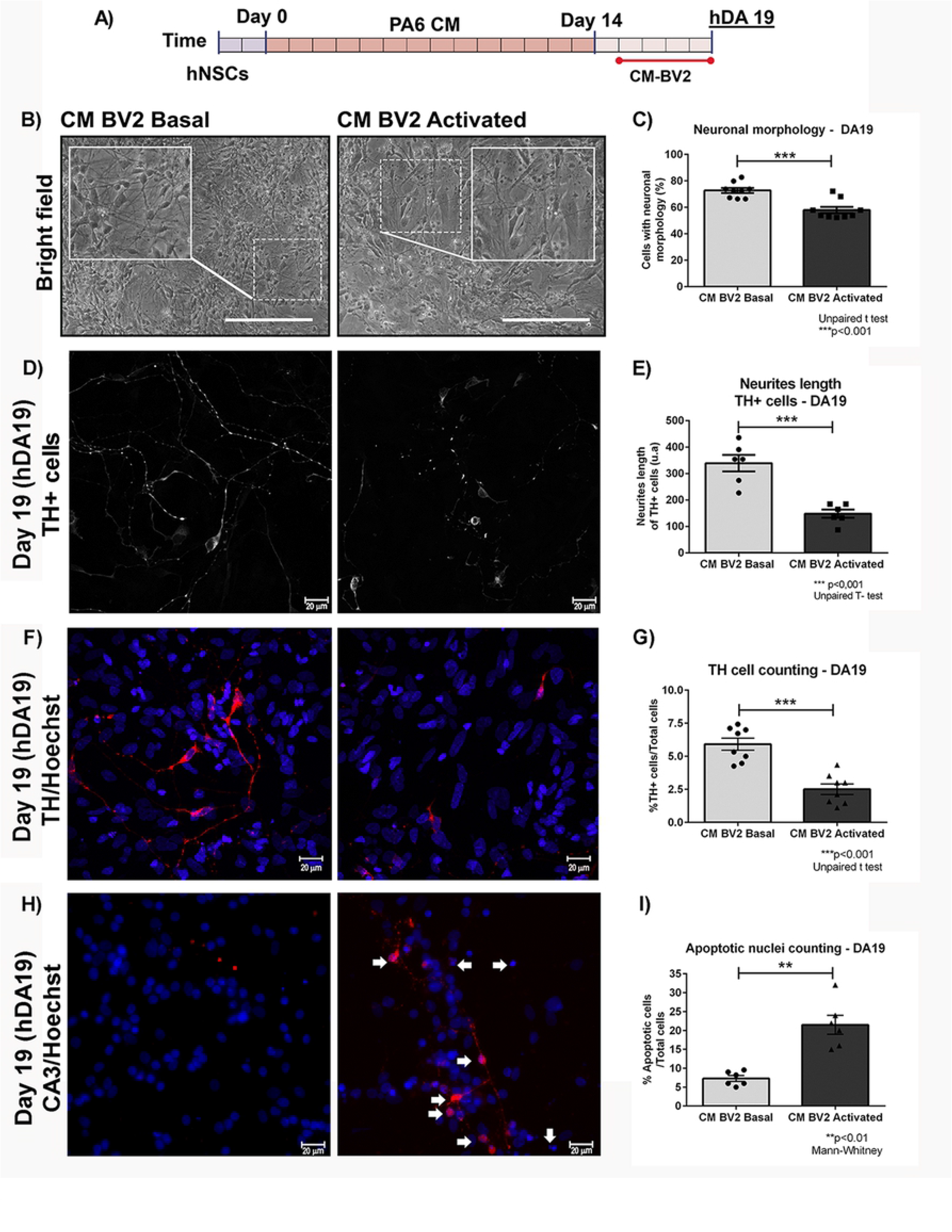
Effect of the activated microglia CM in the differentiation of DA precursors. (A) DA15 cultures were exposed to microglia CM (Basal or Activated) until day 19 (DA19). (B-E) For semi-quantitative analysis, photographs from independent experiments were analyzed to determine neuron-like cell count and neurites length. Asterisks indicate statistically differences (***p<0.001. Unpaired T-test. n=7) in percentage of neural-cell-like (C) and neurite length (E) of DA precursors cultured under inflammatory conditions (CM-BV2 activated) versus basal (CM-BV2 basal). There is a decrease in the number of cells with neuronal morphology and neurite length in DA precursors exposed with CM-BV2 activated. (F) Confocal images of TH-stained DA19 cell cultures. G) Asterisks indicate statistically differences (***p<0.001. Unpaired T-test. n=9) for cell counting assays of TH+ cells of DA precursors cultured with CM-BV2 activated versus basal (CM-BV2 basal). (H-I) Cell death was analyzed by fluorescence microscopy using Hoechst and detection of activated caspase 3 (CA3) by immunofluorescence. (H) Arrowheads indicate apoptotic nucleus and CA3-immunoreactive cell. An increment of CA3-positive cells (see indicator arrows) was observed in DA precursors incubated with CM-BV2 activated (40x magnification). (I) Asterisks show statistically significant differences (**p<0.01. Mann Whitney. n=6) in DA precursors incubated with CM-BV2 activated versus CM-BV2 basal. Values are means±SEM for the percentage of apoptotic cells relative to the total cell number.

### Conditoned media from microglia have toxic effects on hDAp: long term effects

In order to know if the short term effects after acute exposure to inflammatory conditions were transient or not, the protocol described before were carried out until DA19. Then, cell medium was replaced with PA6-CM (DA differentiation medium) and then, at the final stage of DA differentiation protocol (DA28), survival and morphological parameters were studied (Fig. 5A). At DA28, a statistical reduction in the percentage of neuronal-like cells (CM-BV2 Basal: 89,6±2,9 vs. CM-BV2 Act: 64,5±3,9 %. ***p<0,001) and neurite length (CM-BV2 Basal: 383,6±24,9 vs. CM-BV2 Act: 183,5±24,1. ***p<0,001) were detected in those cultures who were exposed for an acute period of time to a microglial activated-environment (CM-BV2 Activated) (Fig. 5B-E). Moreover, a reduction in the percentage of TUJ1-positive cells, in cell cultures exposed with CM-BV2 Activated was obseved (CM-BV2 Basal: 54,0±3,1 vs. CM-BV2 Act: 29,8±0,9 %TUJ1+cells. ***p<0,001) (Fig. 8C, 8E). In addition, our results showed significant decrease in the percentage of TH-positive cells in DA cultures treated with CM from activated BV2 (CM-BV2 Basal: 9,8±0,5 vs. CM-BV2 Act: 4,3±0,5 %TH+cells. ***p<0,001) (Fig. 5F, 5G). We conclude that there is no recovery of TH-positive cells in cultures initially incubated under inflammatory conditions.

**Fig 5.**
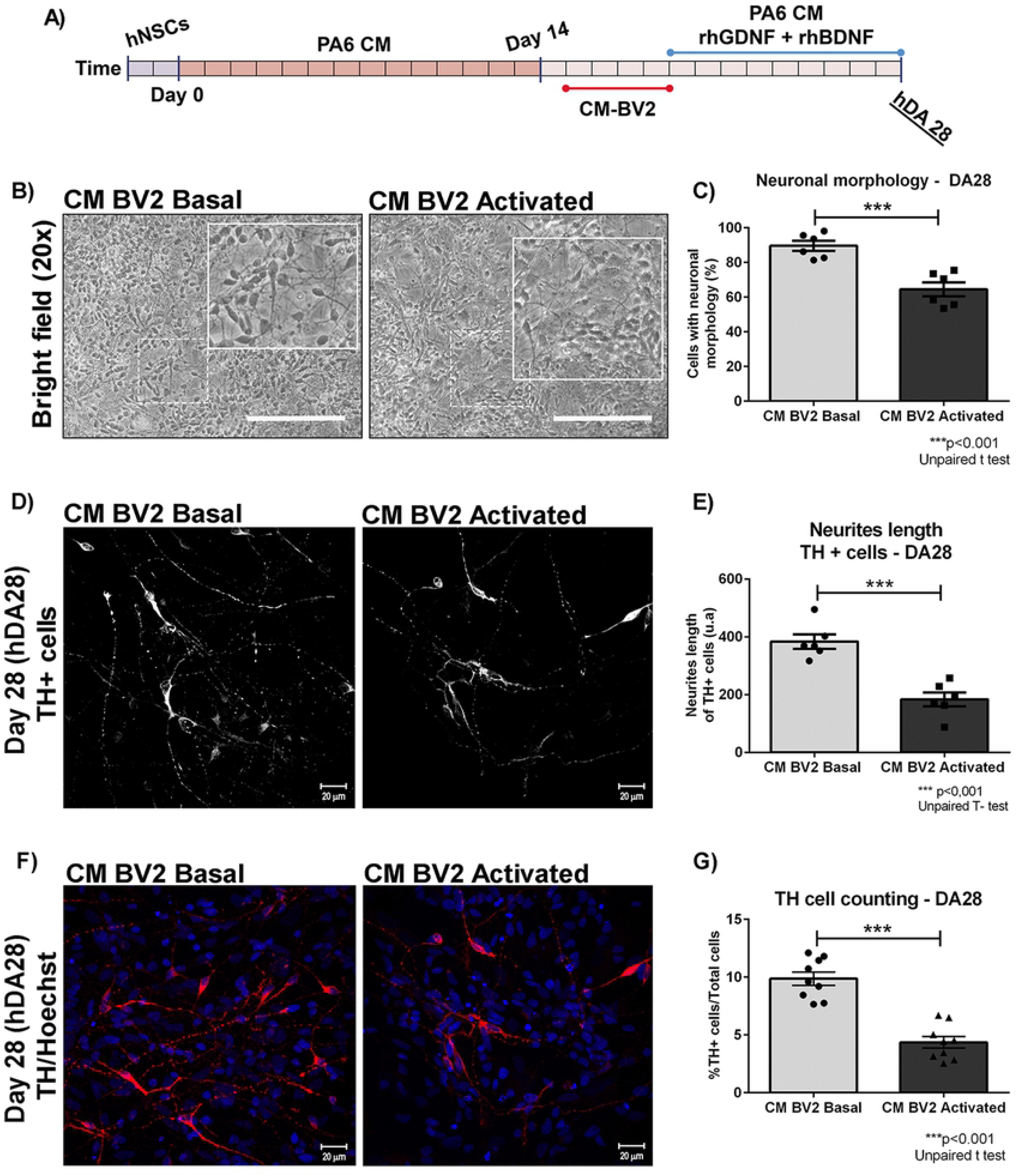
Impact of pro-inflammatory microenvironment on DA cultures. (A) CM from basal and activated condition were added to DA precursors and incubated during 4 days. Morphological analysis and detection of DA neurons were performed at DA28. (B-E) For semi-quantitative analysis, photographs from independent experiments were analyzed to study morphological alterations and determine neuron-like cell count and neurites length. Asterisks indicate statistically significant differences at DA28 stage (***p<0.001. Unpaired T-test. n=6) in the percentage of neural-cell-like (C) and neurite length of TH-positive cells (***p<0.001. Unpaired T-test. n=6) (E) of hDAp cells cultured under inflammatory conditions (CM-BV2 activated) versus basal (CM-BV2 basal). (F-G) Cell counting of TH-positive cells was performed. Representative photomicrographs from TH immunofluorescence (40x) of DA28 cells obtained from DA28 cultured with CM-BV-2 (basal and activated condition) are shown. Asterisks indicate statistically significant differences (***p<0.001. Unpaired T-test. n=6) for cell counting assays of TH+ cells of hDAp cultured under inflammatory conditions (CM-BV2 activated) versus basal microenvironment (CM-BV2 Basal) (G). Values are means±SEM of n independent trials.

### Functional effects of TNF-α on survival and differentiation of DA precursors

Previously, the pro-inflammatory cytokine TNF-α was observed to be expressed after transplantation (Fig. 2G), which was detected in cell culture supernadant of activated microglia (Fig. 3B). Earlier studies reported citotoxic effects of TNF-α in neural cell lines (20). These findings lead us to study the role of TNF-α in the differentiation and survival of hDAp.

DA15 were cultured with CM-BV2 (basal or activated condition) during 4 days as previously described, with or without Etanercept (a TNF-α inhibitor) as co-treatment (Fig. 6A). Neuronal differentiation was analysed by cell morphology and TUJ1-cell counting. Acute exposure to CM from activated microglia (CM-BV2 Act) affected cell morphology and TUJ1-positive cell percentage (CM-BV2 Basal: 43,9±1,4 vs. CM-BV2 Act: 30,3±1,4 %TUJ1+cell. ***p<0,001), while TNF-α blockage was able to reverse neuronal loss (CM-BV2 Act+Etan: 40,9±1,7 vs. CM-BV2 Act: 30,3±1,4 %TUJ1+cell. **p<0,01) (Fig. 6B-E).

**Fig 6.**
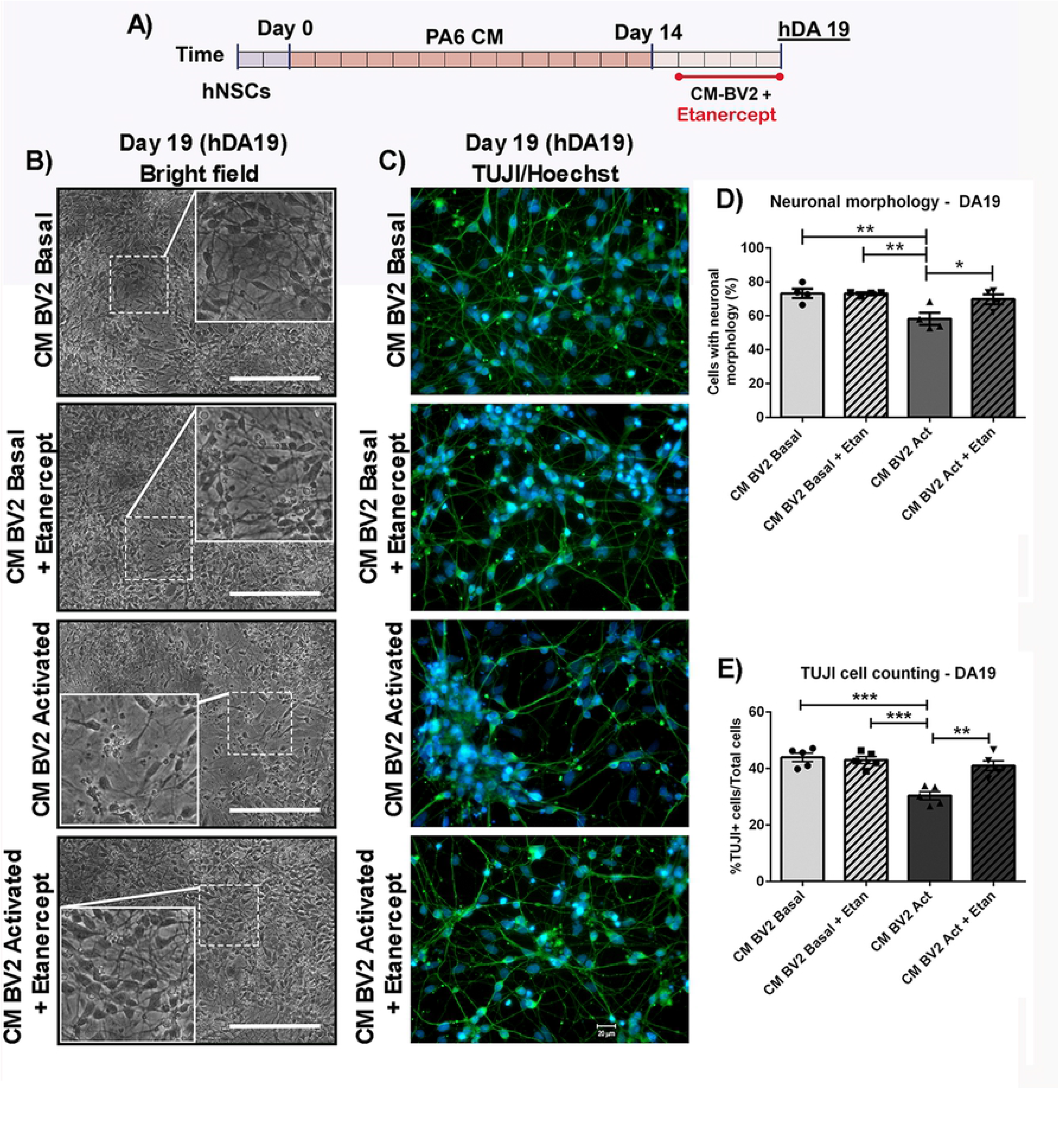
Effect of TNF-α inhibitor on differentiation of DA precursors exposed to activated microglia CM. (A) DA precursors were exposed with CM-BV2 during the 4 days, in the presence of Etanercept (a TNF-α inhibitor). (B) For semi-quantitative analysis, photographs from independent experiments were analyzed to determine neuron-like cell count. (C) Detection of TUJI+ cells by immunofluorescence and cell counting were performed at DA19. Photomicrographs are shown (40x). (D) Asterisks indicate statistically significant differences in percentage of neural-cell like DA precursors cultured under inflammatory conditions (CM-BV2 activated) versus basal (CM-BV2 basal) (*p<0.01 CM-BV2 Act vs. CM-BV2 Basal). Inhibition of TNF-α prevents the decrease in the number of cells with neuronal morphology (*p<0.05 CM-BV2 Act vs. CM-BV2 Act+Etan). ANOVA followed by Tukey’s post hoc test. n=4 independent assays. (E) Asterisks indicate statistically significant differences in percentages of TUJI+ cells of hDAp cultured with CM from activated BV2 cultures versus CM from BV2 cells under basal conditions (***p<0.001 CM-BV2 Act vs. CM-BV2 Basal). Co-incubation of hDAp with Etanercept inhibited TUJI+ cells diminution (**p<0.01 CM-BV2 Act vs. CM-BV2 Act+Etan). ANOVA followed by Bonferroni test. n=5 independent assays. Values are means ± SEM of n independent trials.

In addition, Etanercept had no effects on the expression of TH when hDAp were incubated with CM from basal BV2 cells. However, inflammatory-mediated suppression of TH expression was overtly reversed by Etanercept co-treatment (CM-BV2 Act+Etan: 5,7±0,5 vs. CM-BV2 Act: 2,6±0,5 %TH+cells. **p<0,01) (Fig.7B, 7E). Quantification of neurite length of DA19 indicated that, while pro-inflammatory conditions caused a decrease in neurite length, co-treatment with Etanercept reduced these alterations in TH-positive cells (Fig. 7C, 7D). Finally, we measured cell death in DA19. Our results suggest that TNF-α mediated inhibition partially reduced the percentage of apoptotic cells (Fig. 7F).

**Fig 7.**
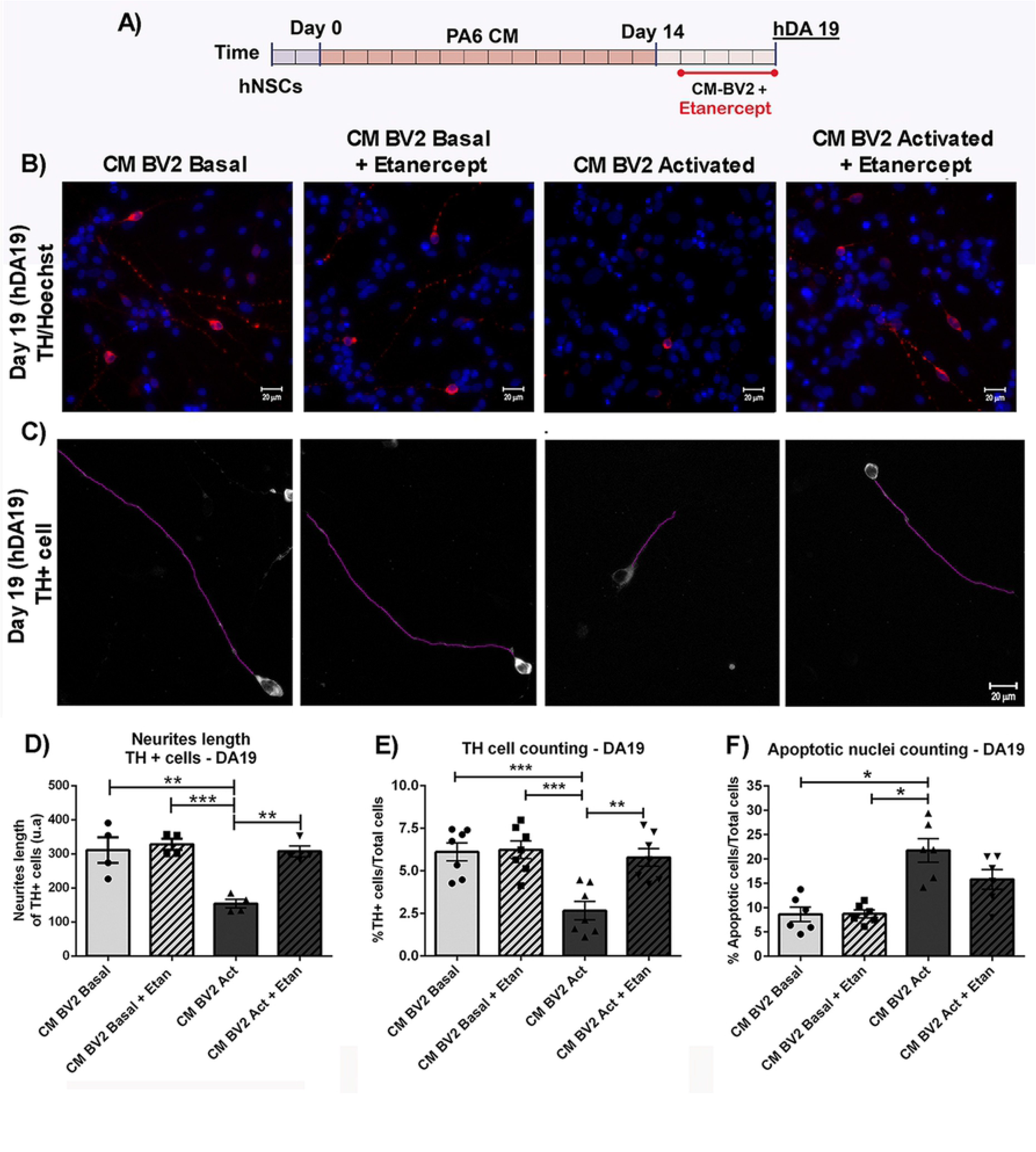
Effect of TNF-α inhibitor on survival and neurite length of DA precursors exposed to inflammatory conditions. (A) DA precursors were cultured in the presence of CM-BV2 (basal or activated condition) during 4 days, in the presence of Etanercept as co-treatment. (B) Detection of TH+cells by immunofluorescence and cell counting were performed at DA19. Photomicrographs from immunofluorescence are shown (40x). (C) Neurite length analyzes of TH+ cells were also performed at DA19. Photomicrographs are shown (40x). (D) Decrease in neurite length were observed after exposure of DA precursors to activated CM-BV2 (**p<0.01 vs. CM-BV2 Basal). Inhibition of TNF-α reduces alterations in neurite length of TH+ cells (**p<0.01 CM-BV2 Act vs. CM-BV2 Act+Etanercept). ANOVA followed by Bonferroni test. n=4. (E) Exposure of DA precursors to activated CM-BV2 decreased the percentage of TH+ cells (*** p <0.001 vs. CM-BV2 Basal). Inhibition of TNF-α prevents TH+ cell loss (**p<0.01 CM-BV2 Act vs. CM-BV2 Act+Etan. ANOVA followed by Bonferroni test. n=7). (F) Cell death was analyzed by apoptotic nucleus counting after Hoechst staining. The results showed that exposure of DA precursors to CM-from activated microglia significantly increase in cell death (*p<0.05 vs. CM-BV2 Basal. Kruskal-Wallis test ANOVA followed by Dunn’s test. n=7). Partial reduction of apoptotic cells are detected in cell culture treated with Etanercept. Values are means±SEM of n independent trials.

Similar effects of TNF-α inhibition were observed in all parameters studied previously at the terminal differentiation stage (DA28), suggesting a persistant protection of the hDAp after Etanercept treatment (Fig. 8 and 9).

**Fig 8.**
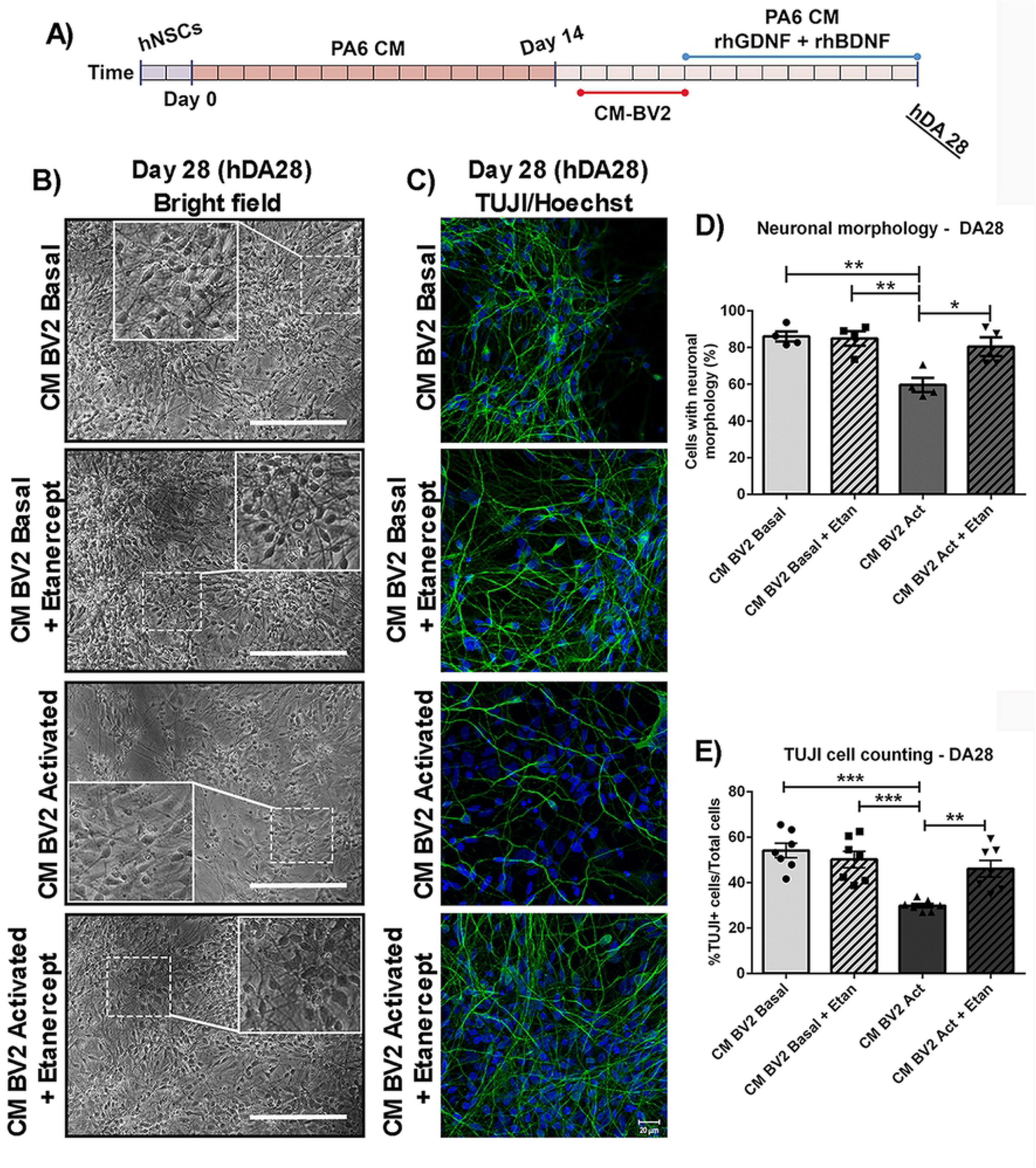
Inhibition of TNF-α and final differentiation of DA precursors exposed to proinflammatory conditions. (A) DA precursors were exposed with CM-BV2 (basal or activated condition) during the 4 days, in the presence of Etanercept. At DA19, cell media was changed to PA6-CM. Morphological assay and detection of TUJI+ cells by immunofluorescence were performed at DA28. (B, D) For semi-quantitative analysis, photographs from independent experiments were analyzed to determine neuron-like cell count. Asterisks indicate statistically significant differences in percentage of neural-cell-like of DA precursors cultured under inflammatory conditions (CM-BV2 activated) versus basal (CM-BV2 basal) (**p<0.01 CM-BV2 Act vs. CM-BV2 Basal). Inhibition of TNF-α prevents the decrease in the number of cells with neuronal morphology (*p<0.05 CM-BV2 Act vs. CM-BV2 Act+Etan). ANOVA followed by Tukey’s post hoc test. n=4. (C, E) Photomicrographs from TUJ1 immunofluorescese (40x) of DA28 cultures are shown. Asterisks indicate statistically significant differences in percentages of TUJ1+ cells of DA cultures exposed with CM from activated BV2 cultures versus CM from BV2 cells under basal conditions (***p<0.001 CM-BV2 Act vs. CM-BV2 Basal). Co-incubation of DA cell cultures with Etanercept inhibited TUJI+ cells diminution (**p<0.01 CM-BV2 Act vs. CM-BV2 Act+Etan.) (n=7 independent assays). ANOVA followed by Bonferroni test. Values are means±SEM of n independent trials.

**Fig 9.**
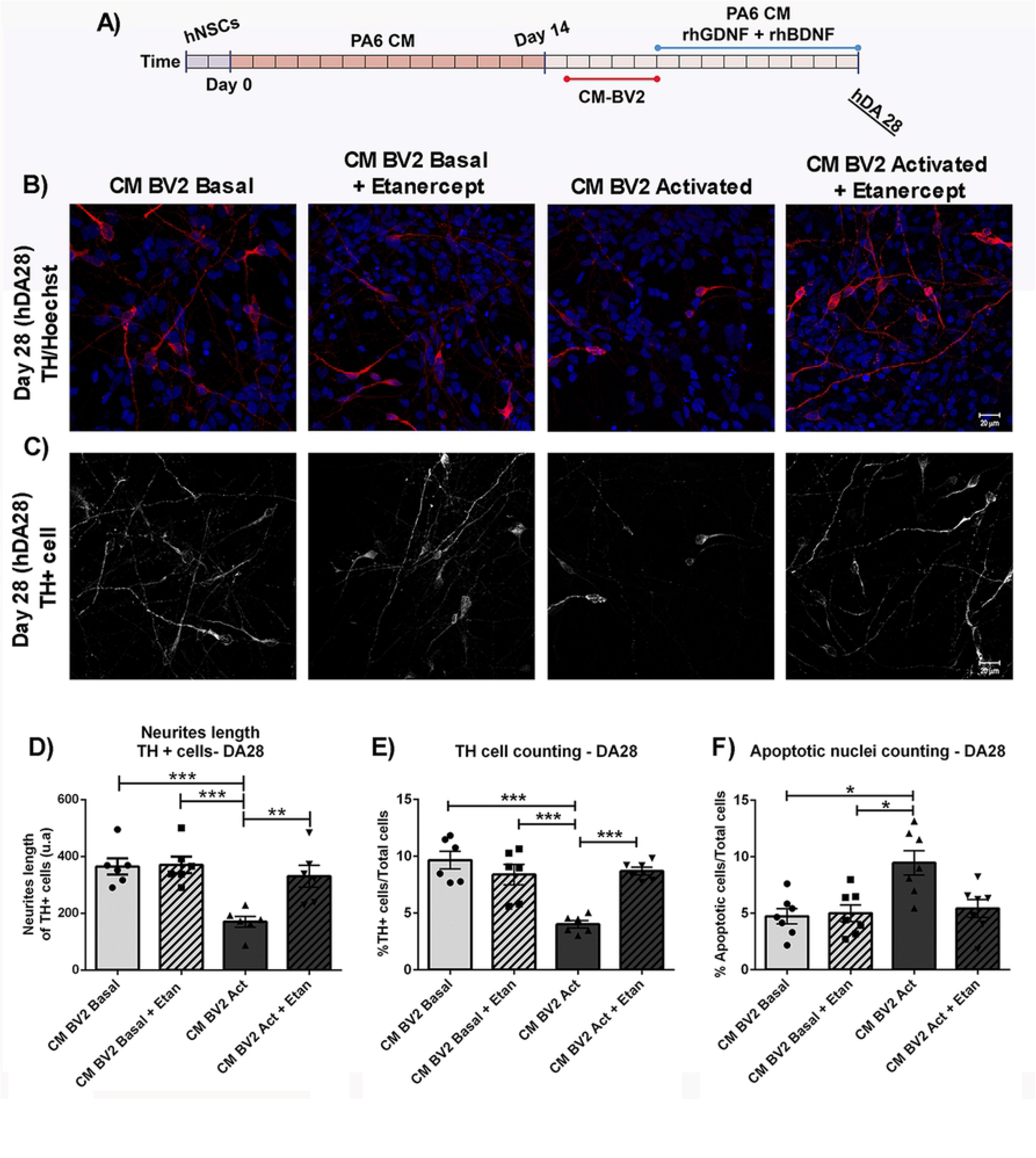
Long term study of Etanercept effect on DA cultures exposed to acute inflammatory conditions. (A) DA precursors were cultured in the presence of CM-BV2 (basal or activated condition) during the 4 days. A co-treatment of DA precursors with CM-BV2 and Etanercept were performed. (B-C) At DA28, immunofluorescence agaisnt the dopaminergic marker TH and neurite length analyzes were performed. Cell death was analyzed by apoptotic nucleus counting after Hoechst staining. Photomicrographs from immunofluorescence are shown (40x). (D) Diminution in neurite length were observed in DA cultures incubated to CM from activated BV2 (***p<0.001 CM-BV2 Act vs. CM-BV2 Basal). Inhibition of TNF-α reduces alterations in neurite length of DA cells (**p<0.01 CM-BV2 Act vs. CM-BV2 Act+Etan. ANOVA followed by Bonferroni test. n=6)). E) Acute exposure of DA precursors to CM from activated microglia decreased the final percentage of TH+ cells (***p<0.001 CM-BV2 Act vs. CM-BV2 Basal). Etanercept co-incubation was able to prevent TH+ cell loss (***p<0.001 CM-BV2 Act vs. CM-BV2 Act+Etan. ANOVA followed by Bonferroni test). (F) The results from cell death analyses show that acute exposure of DA precursors to CM-from activated microglia increase in cell death (*p<0.05 vs. CM-BV2 Basal. Kruskal-Wallis test ANOVA followed by Dunn’s test. n=7). Etanercept was able to reduce the percentage of apoptotic cells. Values are means±SEM of n independent trials.

## Discussion

Cell replacement therapy involves disruption of the blood–brain barrier (BBB) and host tissue damage, which cause astrocyte and microglia activation (21, 22). Therefore, grafted cells are surrounded by an altered environment where host tissue signals could affect relevant processes for the efficacy of cell therapy, such as survival and differentiation of DA precursors.

In this study, we have investigated for the first time the short-term response of the cerebral parenchyma to human DA precursors (hDAp) transplantation and further studied the effects of the microglia response on hDAp viability and differentiation *in vitro*. We show that a glial response was sustained in time after transplantation, together with TNF-α expression, under immunosuppression conditions. *In vitro*, acute exposure to conditioned media (CM) from activated microglia diminished the percentage of TH positive cells, induced cell death and affected the differentiation process. In addition, this acute pro-inflammatory treatment of hDAp had a negative impact on terminal differentiation. Finally, specific inhibition of TNF-α reduced the loss of hDAp and the alterations in morphology.

In our *in vivo* model, a short-term host primary response related to the grafted hDAp (DA14) was detected with a significant increase of host MHCII- and GFAP-positive cells in adult immunosuppresed male rats. These observations were supported using other cell types who demonstrated an early increase in Iba1- and GFAP-positive cells following a NSC graft, until day 3 post-surgery (22). Further support to our observations come from work by Tomov and colleges who observed microglia and GFAP-positive cells between 7 and 28 days after allogeneic transplantation of ventral mesencephalic (VM) cells in a rodent model of PD (6, 23). In addition, MHCII-positive cells around hDAp grafts derived from iPSCs were detected long-term in a PD model of immunosuppressed non-human primates (24). At the molecular level, we observed expression of the pro-inflammatory cytokine TNF-α in host-microglia (ED-1)-positive cells after transplantion with hDAp. Interestingly, TNF-α was also detected on the acute period following VM neuroblasts allogeneic grafts from rodents (25) and in allogeneic and xenogeneic transplantation of VM neuroblasts from rodents and pig, respectively (26). Therefore, our data extend and support previous observations on an early host response after brain grafting of other cell types and animal models. Taken together, we preliminary conclude that there seems to be no overt specificity on the host innate immune response to different transplanted cell types at the cellular level.

We also developed an *in vitro* approach which partially simulates the pro-inflammatory microenvironment from the host response related to the graft. This system is based on exposure of hDAp derived from human NSCs to conditioned media from activated BV-2 microglial cells at short-(DA19) and long-term (DA28) end points. Previous research has demonstrated that BV2 cells are a valid model of primary microglia culture (27). In addition, rodent TNF can activate human TNFRI and TNFRII (28). Other microglial models such as primary microglial cultures or human iPSC-derived microglia could be used in future experiments to test similar hypotheses as in this work. Our results showed a significant increase in cell death of DA precursors exposed with CM from activated microglia by means of a decrease in TH-positive cells in early and late cell culture stages. Our data on cell death extend similar effects of CM from activated microglia observed in other cell types such as SH-SY5Y and PC12 cultures (10, 15). Previous reports suggest that the crosstalk between the Bcl family and NF-κB could be involved in DAn vulnerability (29, 30). The functional role of these molecules required further analyses.

We also observed morphological alterations specifically induced by CM from activated microglia, such as a decrease in neuron-like cells and neurite length of TH-positive cells at both stages of DA differentiation. Moreover, the percentage of TUJ1-positive cells, a pan-neuronal marker, was diminished by microglia activation. Interestingly, using human cortical neural progenitor cells, TNF-α treatment during six days reduced TUJ1 percentage and increase GFAP-positive cells, suggesting that this cytokine inhibited neuronal differentiation (31). Altogether, our results and others indicate that activated microglial cells and TNF-α could play a role in the survival and differentiation of hDAp and other cells after transplantation.

From the evidence obtained *in vivo*, we were interested in analyzing the effect of TNF-α on hDAp. Co-treatment of activated CM with Etanercept, a TNF-α inhibitor, was able to reverse the reduction of TH-positive cells, cell death and morphological alterations previously observed in hDAp. These results extend a previous finding which reported that inhibition of TNF-α reversed the reduction of DA markers and morphological alterations in other cells such as human TH-positive cells derived from Synovial adipose stem cells (32).

As we mentioned above, none of the current PD therapies stops neurodegeneration or functionally replaces dopaminergic neuronal loss. Currently, a remarkable effort is being made in order to take cell replacement therapy for PD to the clinic (5). Recently, in 2018, the first clinical trial using GMP-grade hDAp derived from iPSCs was launched, a case report of autologous-cell therapy for PD was published last year and a phase 1 study to evaluate pluripotent stem cell-derived hDAp in patients with PD was approved by the regulatory authorities of US (33, 34). Survival, differentiation and integration of the transplanted precursors are biological processes that could influence the effectiveness of this strategy and are affected by the host response (21).

In particular, a major limiting factor of cell replacement therapy for PD is still the poor survival rate (10%) of grafted DA precursors (35). Our study points to TNF-α inhibition as a possible strategy to increase survival and differentiation of grafted hDAp. It remains to be determined whether the sole inhibition of TNF or any of its receptors after transplantation is necessary and sufficient to inhibit the deleterious effects of inflammation as we have observed *in vitro*. Nevertheless, several TNF-α inhibitors are clinically available for other diseases. A possible strategy that could be used is the transplantation of DA precursors along with co-infusion of a TNF inhibitor or monoclonal antibodies since inhibitors against TNF-α such as Etanercept and TNF-R1 antagonist as ATROSAB cannot cross BBB under physiological conditions. Alternatively, since cell transplantation include temporal BBB disruption, treatment with agents to neutralize TNF-α deleterious action could be used immediately after surgery. On the other hand, the peripheral administration of a soluble TNF-α inhibitor (XPro1595) was neuroprotective on an *in vivo* model of PD (36). Then, molecules such as XPro1595 could be good candidates to be used in animals models of cell replacement therapy for PD in order to analyze its potential in this specific strategy.

In conclusion, our data indicate that microglia-derived TNF-α plays a key role in the possible effects of the host response to hDAp transplantation by affecting survival and differentiaton at short and long-term. Selective targeting of TNF-α holds translational potential to increase survival and differentiation of DA precursors even under immune suppressive treatments targetting the adaptive immune response. Finally, the *in vitro* model described might be useful to study the mechanism of action of microglia on hDAp and search for potentials anti-inflammatory and/or neuroprotective treatments in order to improve survival and differentiation efficacy of hDAp.

## Author Contributions

Conceptualization: FJP. Investigation and Methodology: SDW, VG, CG, MIF, JB. Formal Analysis: SDW, FJP. Project Administration: SDW, FJP. Funding Acquisition: FJP, SDW. Writing – Original Draft Preparation: SDW, FJP. Writing – Review & Editing: CF, CG. SDW, CF, CG, JB, FJP are members of the research career of CONICET. MIF is member of the MTA career of CONICET.

## Notes

### Competing Interest Statement

The authors have declared no competing interest.

